# A systematic exploration of unexploited genes for oxidative stress in Parkinson’s disease

**DOI:** 10.1101/2024.03.11.583425

**Authors:** Takayuki Suzuki, Hidemasa Bono

## Abstract

Human disease-associated gene data are accessible through databases, including the Open Targets Platform, DisGeNET, miRTex, RNADisease, and PubChem. However, missing data entries in such databases are anticipated because of factors, such as errors/biases by curators and text mining failures. Additionally, the extensive research on human diseases has resulted in challenges to register comprehensive data. The lack of essential data in databases hinders knowledge sharing and should be addressed. Therefore, we propose an analysis pipeline to explore missing entries of unexploited genes in the human disease-associated gene databases. To demonstrate this, we used the pipeline for genes in Parkinson’s disease with oxidative stress, which revealed two unexploited genes: nuclear protein 1 (*NUPR1*) and ubiquitin-like with PHD and ring finger domains 2 (*UHRF2*). The proposed methodology and findings facilitate the identification of disease-associated genes that are not completely represented in existing databases, thereby facilitating easier access to the potential human disease-related functional genes.

## INTRODUCTION

Human disease research is a significant area in biology. For example, querying “parkinson disease”[All Fields] in the PubMed literature database yields 94,062 literatures (February 11, 2024). Subsequently, research findings are curated by experts or extracted through text-mining methodologies to register in databases that facilitate collective intelligence. For instance, the Clinical Genome Resource (ClinGen)^1^, developed by the National Institutes of Health (NIH), curates, assesses, and disseminates aggregated genetic and disease associations as public data. The GWAS Catalog^2^ compiles genome-wide association data, including single nucleotide polymorphisms (SNPs) with associated disease risks. Open Targets Platform^3^ integrates data from ClinGen and GWAS Catalog, and other public human disease-related databases, including CRISPRBrain^4^, Open Targets Genetics^5^, Gene Burden^6,7^, and Europe PMC^8,9^. Additional databases, including DisGeNET (expert-curated integrated gene-disease associations until June 2020)^10^, miRTex (specialized in text mining approach-extracted microRNA)^11^, RNADisease (specialized in collecting non-coding RNAs with integrated approaches of curating and text-mining)^12^, and PubChem (specialized in extracted data from PubMed)^13^ offer complementary insights. They use various combinations of integration of data, specific text mining methods, and focus on RNA-disease links, capturing disease-gene associations that may not be included in the Open Targets Platform. These five databases offer comprehensive access to the latest human genes that have been implicated as functionally related to the disease.

As previously mentioned, PubMed contained 94,062 studies related to PD. With these publications and accompanying data, missing data entries for disease-related functional genes are anticipated in databases^14^. Factors that contribute to missing disease-related genes may include challenges in computationally accessing biomedical statement contexts and supplementary data in the literature, oversights or biases by curators, and text-mining failures. These unexploited genes, which can be referred to as false-negative genes in current gene-disease association databases, represent incomplete dissemination of prior knowledge, potentially hindering research advancement and the development of disease prevention and treatment strategies. The manual identification of missing data entries across the literature databases requires substantial human resources and time expenditures. Therefore, we propose an approach to identify unexploited genes, which are missing data entries in five relevant databases. Based on our previous research on oxidative stress (OS)^15,16^, we selected PD, which exhibits pathological associations with OS, to demonstrate the efficacy of our methodology for identifying unexploited genes as proof of concept.

PD is a neurodegenerative disorder affecting over 6 million patients worldwide^17^. The primary symptoms observed in patients with PD include unilateral rigidity, bradykinesia, tremors, and non-motor symptoms, such as cognitive dysfunction^18–20^. The defining characteristics of PD include disordered α-synuclein aggregation, Lewy body formation, and significant loss of dopaminergic (DA) neurons in the substantia nigra, resulting in depleted dopamine levels, causing motor and cognitive deficits^18,20–23^. PD has been considered to be closely associated with the biological phenomenon, oxidative stress (OS), wherein reactive oxygen species (ROS) and nucleophiles both contribute to and are generated by aggregating α-synuclein, Lewy bodies, and DA neuron loss^24^. Because OS is characterized by an imbalance between ROS levels and antioxidant defenses, substantial evidence has implicated it in PD pathogenesis^24^. As current therapies focus on symptomatic treatment, such as dopamine replacement, rather than root-cause therapy, understanding the molecular mechanisms of PD symptoms associated with OS is crucial for developing more effective therapies or biomarkers^25^. Querying “parkinson disease”[All Fields] AND “oxidative stress”[All Fields] in PubMed yields 4,061 publications (as of February 15, 2024), and manually reviewing all 4,061 articles to sequentially identify unexploited genes would be labor-intensive. Therefore, the discovery of unexploited genes as candidates for the field of PD and OS research is necessary to further understand their underlying mechanisms.

To identify unexploited genes, we developed an analytical pipeline consisting of four significant stages. First, we curated a candidate gene set based on disease-relevant gene expression data. Second, we classified candidate genes based on the presence or absence of disease associations in the five applicable databases. Third, we refined the genes likely to be functional, leveraging data from transcriptome meta-analysis and transcriptome-wide association studies (TWAS). Finally, we manually searched the refined gene list to identify the unexploited genes with documented disease associations in the literature, but no links in the five databases. To identify genes related to OS in PD, we discovered two unexploited PD genes: nuclear protein 1 (*NUPR1*), and ubiquitin-like with PHD and ring finger domains 2 (*UHRF2*). The proposed approach and its findings will facilitate the identification of unexploited genes missing from databases, thereby advancing future research on human diseases.

## RESULTS

### Overview of the pipeline (Figure 1)

Our stepwise methodology entailed the following: 1) Transcriptomic meta-analysis and literature mining were independently conducted to identify differentially expressed genes (DEGs) associated with oxidative stress (OS) and Parkinson’s disease (PD) in the human brain [Figure 1a]. These outputs were compared to extract the dysregulated genes in both OS and PD (OS-PD-DEGs, n=168) [Figure 1a]. 2) The 168 candidate genes in OS-PD-DEGs were categorized into two subsets based on their associations with PD according to relevant databases (Open Targets Platform^3^, DisGeNET^10^, miRTex^11^, RNADisease^12^, and PubChem^13^)—116 genes were not associated with PD (PD-unlinked-genes) [Figure 1b], whereas the remaining 52 genes exhibited PD associations (PD-linked-genes). 3) To identify genes with functions, we filtered PD-unlinked-genes using data from the PD transcriptome meta-analysis and TWAS, with 10 genes (unexploited candidate genes) remaining for the last step. 4) Finally, two unexploited genes *(NUPR1* and *UHRF2*) were discovered by manually searching PubMed Central for data on unexploited candidate genes [Figure 1c].

**Figure 1.**
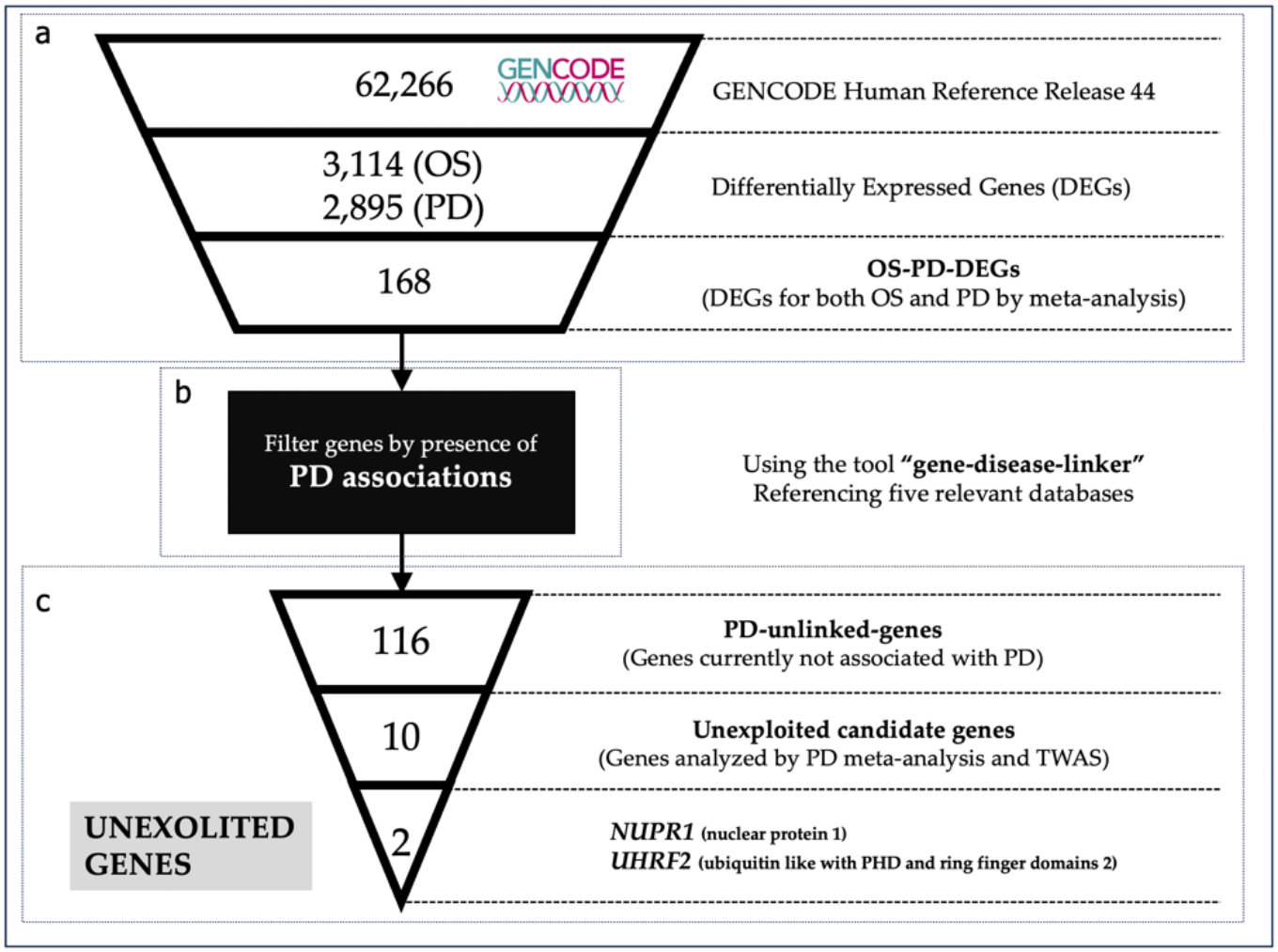
Overview of the pipeline filtering the number of candidates to search unexploited genes. Figure 1a: Transcriptome meta-analysis to retrieve differentially expressed genes (DEGs) in both oxidative stress (OS) and Parkinson’s disease (PD) (n=168). Figure 1b: Gene-disease-linker (see Methods) filters the 168 candidate genes into PD-unlinked-genes based on gene-PD association with evidence studies. Figure 1c: PD-unlinked-genes are further filtered using transcriptome-wide association studies (TWAS) and PD meta-analysis results to obtain unexploited candidate genes. Manual search of unexploited candidate genes revealed two unexploited genes.

### Collection of DEGs through OS

A transcriptomic meta-analysis was conducted on 122 paired RNA sequencing (RNA-seq) datasets^26– 33^ from cultured human cells related to brain to identify DEGs associated with OS. Each pair comprised an OS sample and a matched normal condition sample from the same original study. Specifically, the transcriptomes of neurons, astrocytes, and neural progenitor cells [Supplementary Table1]^34^ under OS or normal conditions were compiled from the Gene Expression Omnibus (GEO) database^35^. Oxidative stressors included radiation, hydrogen peroxide, rotenone, 1-methyl-4-phenylpyridinium (MPP^+^), paraquat, 6-hydroxydopamine (oxidopamine), methyl mercury chloride, and zinc (Table 1) [Supplementary Table1]^34^. A total of 3,114 genes (5% of all genes whose expression was quantified, termed OS-DEGs), were collected as DEGs, consisting of 1,557 most upregulated and downregulated genes [Figure1a) with an ON_score of 1.5. Of the 357 genes, 37 possessed Ensembl IDs annotated to “GO:0006979: response to oxidative stress” were included in OS-DEGs (p-value: 2.671*e^-5).

**Table 1:** Metadata of curated data for meta-analysis of RNA-seq with oxidative stress in brain. Table1a: Number of data pairs retrieved for each source of oxidative stress (OS). Table1b: Number of data pairs retrieved for each type of cell culture.

| a |  | b |  |
| --- | --- | --- | --- |
| Source of Oxidative Stress | Number of Data Pair | Types of cultured cells | Number of Data Pair |
| 1-methyl-4-phenylpyridinium (MPP) | 35 | Neuron | 83 |
| Rotenone | 19 | Astrocyte | 24 |
| H<sub>2</sub>O<sub>2</sub> | 17 | Neural progenitor | 15 |
| Radiation | 15 | Total | 122 |
| Paraquat | 9 |  |  |
| 6-Hydroxydopamine | 9 |  |  |
| Methyl mercury Chloride | 9 |  |  |
| Zinc bis | 9 |  |  |
| Total | 122 |  |  |

### Collection of DEGs through PD

Because existing meta-analyses that have delineated DEGs associated with PD, we curated and compiled PD-DEG sets from three relevant studies^36–38^ (Table 2). These studies performed meta-analyses of PD transcriptomic data to derive robust gene expression signatures for the disease. Additionally, PD-associated genes identified through TWAS in the brain tissue were obtained from the TWAS-Atlas database^39^ and incorporated. In total, the PD-DEG comprised 2,895 unique genes (1,378 from PMID:27611585^36^, 585 from PMID: 33390883^37^, 989 from PMID: 37347276^38^, and 196 from the TWAS-Atlas) [Figure 2]. Gene set enrichment analysis (GSEA) confirmed the signature of the PD-DEG compilation, with the most enriched term as “has05012: Parkinson disease” [Figure 3a]. Collating PD-DEG lists from multiple large-scale omics studies and databases generated an extensive catalog of genes dysregulated in PD for subsequent analyses.

**Table 2:**
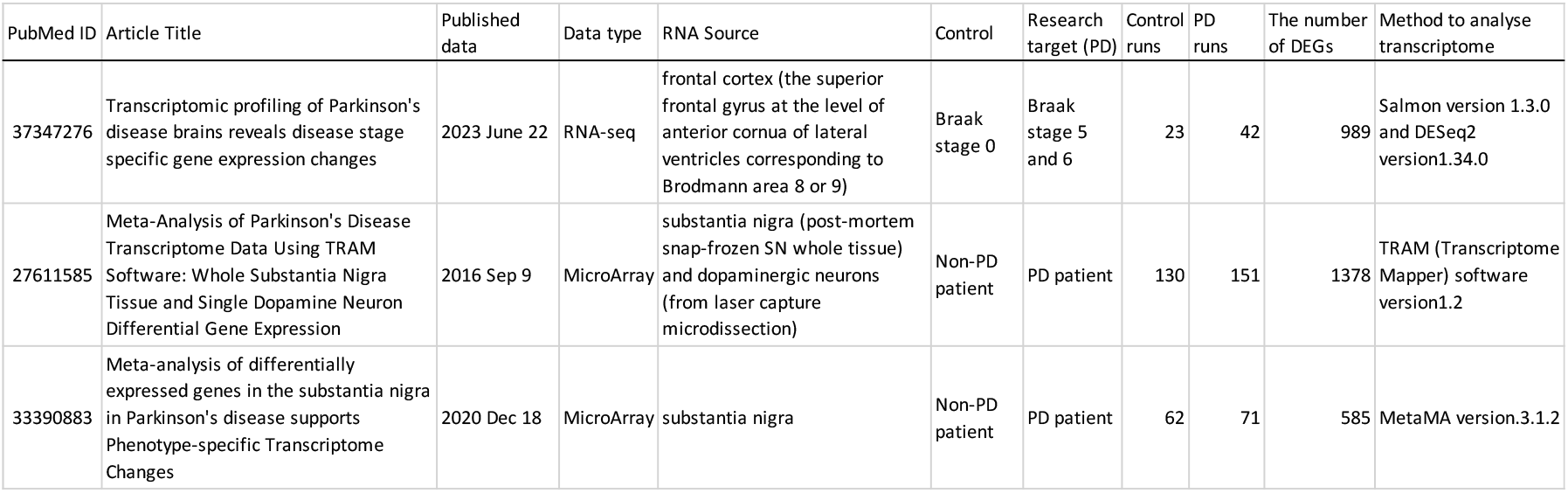
Metadata for curated data for meta-analysis of Parkinson’s disease (PD).

| PubMed ID | Article Title | Published data | Data type | RNA Source | Control | Research target (PD) | Control runs | PD runs | The number of DEGs | Method to analyse transcriptome |
| --- | --- | --- | --- | --- | --- | --- | --- | --- | --- | --- |
| 37347276 | Transcriptomic profiling of Parkinson's disease brains reveals disease stage specific gene expression changes | 2023 June 22 | RNA-seq | frontal cortex (the superior frontal gyrus at the level of anterior cornua of lateral ventricles corresponding to Brodmann area 8 or 9) | Braak stage 0 | Braak stage 5 and 6 | 23 | 42 | 989 | Salmon version 1.3.0 and DESeq2 version1.34.0 |
| 27611585 | Meta-Analysis of Parkinson's Disease Transcriptome Data Using TRAM Software: Whole Substantia Nigra Tissue and Single Dopamine Neuron Differential Gene Expression | 2016 Sep 9 | MicroArray | substantia nigra (post-mortem snap-frozen SN whole tissue) and dopaminergic neurons (from laser capture microdissection) | Non-PD patient | PD patient | 130 | 151 | 1378 | TRAM (Transcriptome Mapper) software version1.2 |
| 33390883 | Meta-analysis of differentially expressed genes in the substantia nigra in Parkinson's disease supports Phenotype-specific Transcriptome Changes | 2020 Dec 18 | MicroArray | substantia nigra | Non-PD patient | PD patient | 62 | 71 | 585 | MetaMA version.3.1.2 |

**Figure 2:**
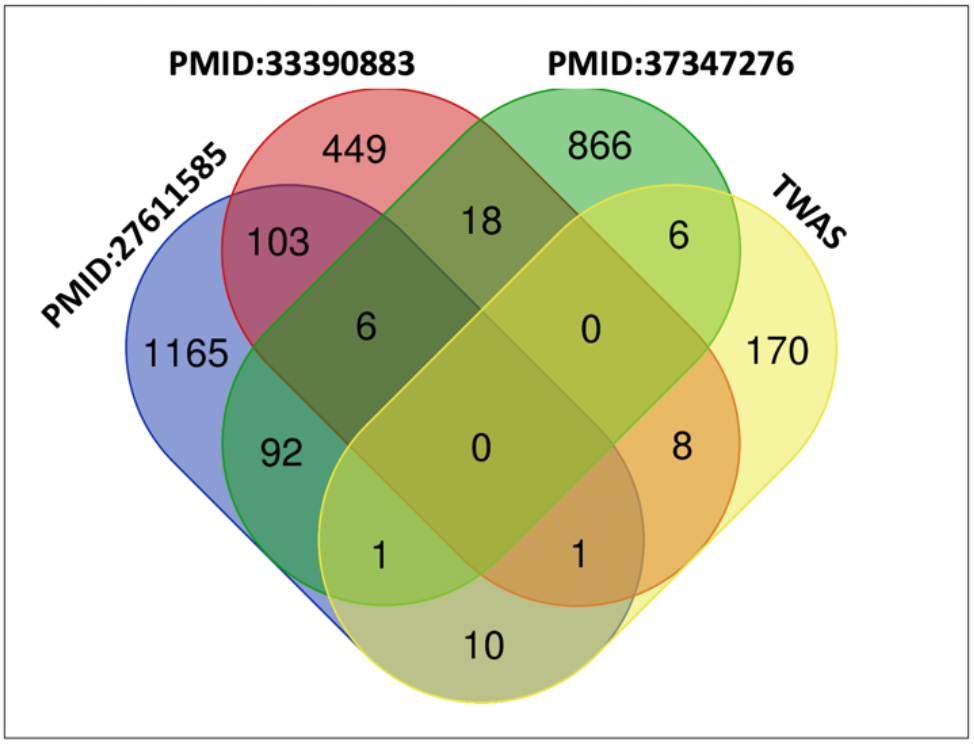
Venn diagram visualizing overlaps of PD-DEGs (n=2,895) among the studies. PMID represents PubMed ID. Transcriptome-wide association studies (TWAS) represents gene sets acquired from the TWAS-Atlas.

**Figure 3a:**
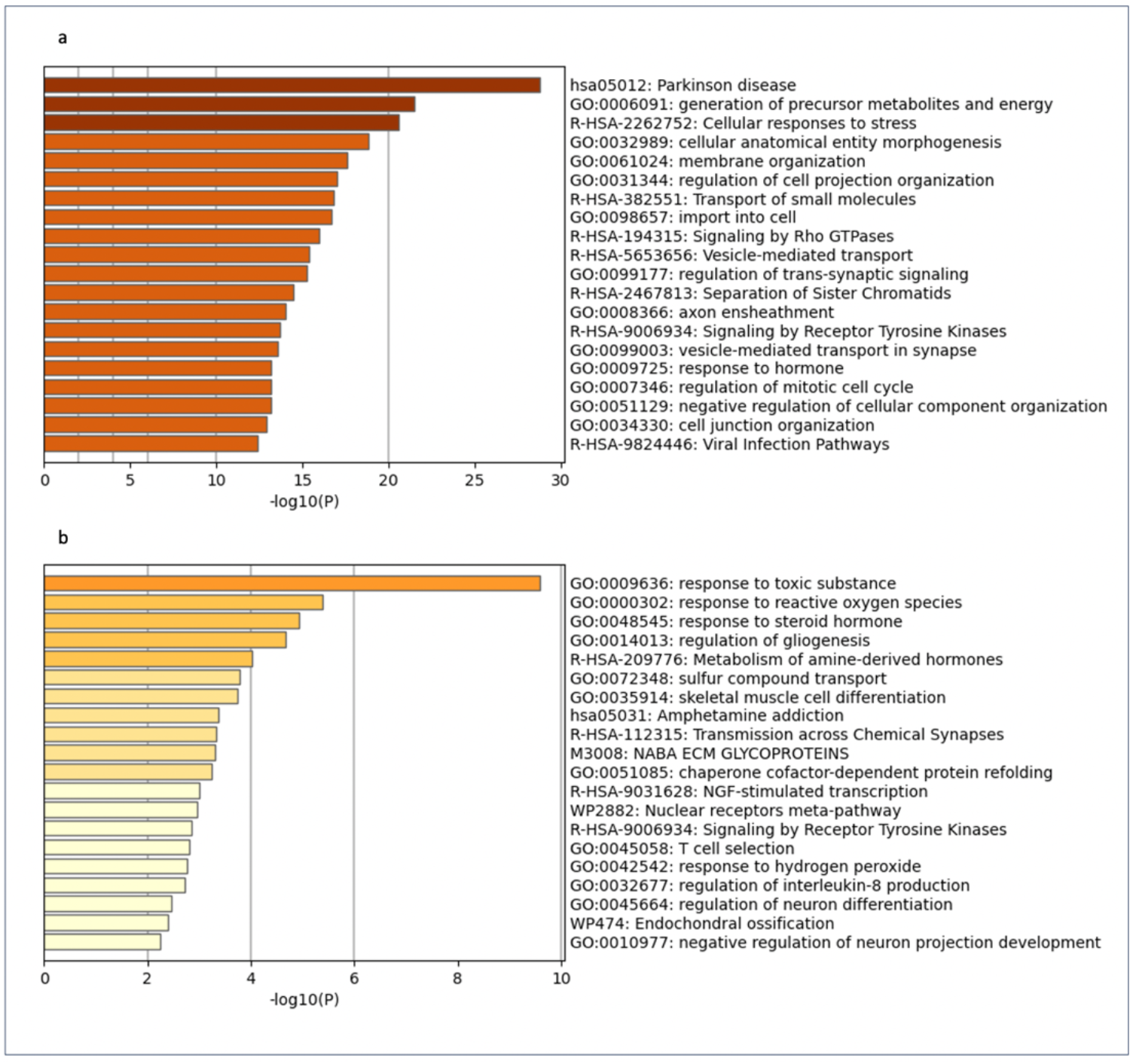
Results of gene set enrichment analysis (GSEA) of the PD-DEGs (n=2,763, unrecognized genes by Metascape, which are unstudied genes lacking NCBI gene IDs). Figure 3b: Results of GSEA of the differentially expressed genes (DEGs) commonly dysregulated in both oxidative stress (OS) and Parkinson’s disease (PD) (n=168).

### Collection of DEGs in both PD and OS

Comparative analysis of the 3,114 oxidative stress differentially expressed genes (OS-DEGs) and 2,895 Parkinson’s disease (PD-DEGs) revealed 168 genes dysregulated in both conditions (termed as OS-PD-DEGs) [Figure 1a). Of these, 132 were protein-coding and 36 were non-coding RNA/pseudogene/small nucleolar RNA or unknown. GSEA of the 168-DEGs overlapping genes exhibited significant enrichment for “GO:0009636: response to toxic substance” and “GO:0000302: response to reactive oxygen species” [Figure 3b), confirming their relevance to these biological processes. Mining of PD associations using the gene-disease-linker^40^ tool identified 52 OS-PD-DEGs with previously reported PD associations (PD-linked-genes). The remaining 116 genes lacking known PD connections were termed as PD-unlinked-genes [Figure 1c]. The results of the gene-disease-linker using OS-PD-DEGs are listed in the Supplementary Table 2^34^. The columns in Supplementary Table2 are described as follows: ENSG: Ensemble gene ID, PD_log2(fold change)_PMID: Log_2_ fold change based on the original papers with PMID specified in the column name, TWAS: whether a gene was indicated in the TWAS, availability of associations: indicates if the gene exhibits association with PD (yes) or not (no), Evidence: databases providing evidence for an association between the given gene and the PD, NU_PMIDs_PD: Number of studies indicating an association between the given gene and PD, NU_PMIDs_NCBI: Number of studies associated with the given gene based on gene2pubmed^41^, PMIDs_PD: PubMed IDs for studies indicating an association between the given gene and PD, and PMIDs_NCBI: PubMed IDs for studies associated with the given gene based on gene2pubmed.

### Analysis of PD-unlinked-genes

Among the 116 PD-unlinked genes, six were identified with TWAS Z-scores in the supplementary data of two studies with TWAS (Table 3, column: original TWAS). Five of these genes were part of a set of 711 genes suggested to confer PD risk in the supplementary data of PMID: 33523105^42^. However, the main texts of the study (PMID: 33523105) lacked mentions of these five genes. *MEI1* was one of the 44 genes implicated in dorsolateral prefrontal cortex PD associations in the TWAS from PMID: 30824768^43^ [Figure 2b and supplementary data], again lacking textual descriptions of *MEI1* within the study (PMID: 30824768). As these six genes were indicated to contribute to PD based on the TWAS results, we presumed them as suitable candidates for assessing the evidence or implications of their involvement in PD molecular mechanisms in the subsequent study mining step of our pipeline. Among the three PD gene expression meta-analyses examined, seven genes were suggested as PD-DEGs in more than two studies (Table 3, column: PD_log2(fold change)). Notably, *NUPR1*, in addition to appearing in the supplementary data of the TWAS-based study, was also selected as a PD-DEG in two PD gene expression meta-analyses, indicating dysregulated expression of *NUPR1* in PD across three studies. Additionally, these seven genes were unexploited candidate genes in the subsequent step. In total of 10 genes [Table 3] have not been previously mentioned their molecular association with PD as textual description in the main texts of studies; however, the sequence data indicated an association. These 10 genes were selected as unexploited candidate genes for the subsequent step, searching the full-text literatures for molecular mechanistic evidence or hypotheses associated with them.

**[Table 3].** Metadata for 10 unexploited candidate genes.

| ensembl gene ID | GeneSymbol | ONscore | PD_log2(fold change)<br>(PMID:37347276) | PD log2(fold change)<br>PMID:27611585 | PD log2(fold_change)<br>PMID:33390883 | TWAS<br>Z score | TWAS original<br>paper (PMID) |
| --- | --- | --- | --- | --- | --- | --- | --- |
| ENSG00000099769 | IGFALS | 27 | Not Found | Not Found | Not Found | 4.45 | 33523105 |
| ENSG00000125703 | ATG4C | 27 | 0.28 | 0.59 | Not Found | Not Found | Not Found |
| ENSG00000112893 | MAN2A1 | 26 | 0.49 | 0.58 | Not Found | Not Found | Not Found |
| ENSG00000147854 | UHRF2 | 25 | 0.29 | 0.58 | Not Found | Not Found | Not Found |
| ENSG00000103932 | RPAP1 | 24 | Not Found | Not Found | Not Found | -5.04 | 33523105 |
| ENSG00000167077 | MEI1 | 24 | Not Found | Not Found | Not Found | -3.37 | 30824768 |
| ENSG00000267454 | ZNF582-DT | 23 | Not Found | Not Found | Not Found | -3.61 | 33523105 |
| ENSG00000176046 | NUPR1 | 23 | Not Found | 0.73 | 0.61 | 4.36 | 33523105 |
| ENSG00000184545 | DUSP8 | 23 | Not Found | Not Found | Not Found | 4.20 | 33523105 |
| ENSG00000164124 | TMEM144 | 21 | 0.91 | -0.83 | Not Found | Not Found | Not Found |

### Identification of unexploited genes

We searched the biomedical full-text literature database PubMed Central with the query “GENE_NAME[All Fields] AND parkinson[All Fields]” to look for unexploited genes (See Methods). Among the 10 genes examined, *NUPR1*, and *UHRF2* were identified as unexploited based on statements in the literature indicating their involvement in PD molecular mechanisms [Table 4]. For *NUPR1*, the PMCID: PMC10734959^44^ was used to determine the gene as unexploited. *NUPR1* was identified as one of the top five as ferroptosis-related hub genes in PD by the methodology using random forest and support vector machines models. Additionally, the association between NUPR1 and alterations of the immune microenvironment of PD patients was indicated by a correlation analysis of NUPR1 and immune characteristics. It was mentioned that “The present study also suggests that *NUPR1* is involved in PD, is positively correlated with PD, and is most likely involved in PD pathogenic mechanisms through ferroptosis and OS”. For *UHRF2*, the PMCID: PMC9775085^45^ was used to determine the gene as unexploited. This review article integrates prior knowledge and proposes that UHRF2 dysregulation contributes to PD progression. It was specifically mentioned as follows; “Altogether, it could be assumed that the dysregulation of *CPNE8, CADPS2*, or *UHRF2* contributes to PD progression via ERK activation induced by the *LRRK2* G2019S mutation”. Table 4 provides the PubMed Central query dates by the author, query results, evidentiary publications (PubMed Central IDs), and quoted text supporting the unexploited status of each gene (evidence statements in the research paper).

**Table 4:** Results of searching unexploited genes. *NUPR1*, and *UHRF2* were identified.

| ensembl gene ID | GeneSymbol | Unexploited genes (yes or no) | Evidence (PMCID) | Evidence statements in research paper | search query in PMC | search date | The number of publications yielded |
| --- | --- | --- | --- | --- | --- | --- | --- |
| ENSG00000125703 | ATG4C | no | NotFound | NotFound | ATG4C[All Fields] AND parkinson[All Fields] | 2024Feb12 | 67 |
| ENSG00000099769 | IGFALS | no | NotFound | NotFound | IGFALS[All Fields] AND parkinson[All Fields] | 2024Feb12 | 13 |
| ENSG00000112893 | MAN2A1 | no | NotFound | NotFound | MAN2A1[All Fields] AND parkinson[All Fields] | 2024Feb12 | 15 |
| ENSG00000147854 | UHRF2 | yes | PMC9775085 | "The downregulation of U | UHRF2[All Fields] AND parkinson[All Fields] | 2024Feb12 | 22 |
| ENSG00000103932 | RPAP1 | no | NotFound | NotFound | RPAP1[All Fields] AND parkinson[All Fields] | 2024Feb12 | 4 |
| ENSG00000167077 | MEI1 | no | NotFound | NotFound | MEI1[All Fields] AND parkinson[All Fields] | 2024Feb12 | 12 |
| ENSG00000176046 | NUPR1 | yes | PMC10734959 | "The top 5 genes were ext | NUPR1[All Fields] AND parkinson[All Fields] | 2024Feb12 | 64 |
| ENSG00000184545 | DUSP8 | no | NotFound | NotFound | DUSP8[All Fields] AND parkinson[All Fields] | 2024Feb12 | 7 |
| ENSG00000267454 | ZNF582-DT | no | NotFound | NotFound | ZNF582-DT[All Fields] AND parkinson[All Fields] | 2024Feb12 | 0 |
| ENSG00000164124 | TMEM144 | no | NotFound | NotFound | TMEM144[All Fields] AND parkinson[All Fields] | 2024Feb12 | 3 |

## DISCUSSION

In this study, we developed a pipeline to identify unexploited genes, that correspond to false-negative genes for a given disease against five databases that provide gene-disease associations. Additionally, it was used to treat PD associated with OS. Through integrated analysis of curated datasets including transcriptomics and transcriptome-wide association results, we filtered the 62,266 genes down to 168 genes (OS-PD-DEGs) that exhibited dysregulation of gene expression in both PD and OS contexts. We subsequently classified OS-PD-DEGs into PD-unlinked and PD-linked-genes based on existing evidence of their involvement in PD. We have further narrowed down the PD-unlinked-genes to 10 unexploited candidate genes. Following a manual search of these 10 genes, *NUPR1*, and *UHRF2* were identified as unexploited genes that were absent from the current gene-disease associations databases. This pipeline effectively identifies unexploited genes by comparing information from the five databases with gene expression data regarding disease-gene associations.

Although it is challenging to conclusively determine the reasons for missing these entries from databases, three potential explanatory factors have been hypothesized as outlined in the Introduction. First, the databases may not have been recently updated. Two evidentiary publications on unexploited genes were recent (published in 2022 and 2023), Therefore, absence of registrations may be possible without updates. Although the Open Targets Platform notes bi-monthly updates, the source database update frequencies vary, potentially explaining the omissions. Second, text-mining extraction failures are possible. For example, text-mining from the Europe PMC relies on co-occurrences at the sentence level with several filtering rules to reduce noise.^9^ Their extraction methodology excludes articles other than research articles and filters out associations that appear only once in the body of an article but not in the article’s title or abstract. Therefore, the *UHRF2*-PD association was excluded from the Open Targets Platform because the review article PMC9775085 was filtered out by the extraction system. Third, certain databases rely on expert manual curation for new data entry, which may be pending or induce human-errors or biases. Using our analytical pipeline to identify unexploited genes helps mitigate these database limitations.

Additionally, the two unexploited genes identified were associated with OS. NUPR1 acts as a key inhibitor of ferroptosis by regulating lipocalin-2 (LCN2) expression to reduce iron accumulation and subsequent oxidative damage^46^. For *UHRF2*, a gene set involved in ROS, UV response, and oxidative phosphorylation was induced in the retinal tissue of Uhrf2 deficient mice^47^. Among the 52 PD-linked genes, the majority have established associations with PD and OS (for instances, *SLC18A2*^48,49^, *TXNIP*^50^, *NEFL*^51,52^, *MPO*^53–56^, *LINC00938*^57,58^). Therefore, the 168 OS-PD-DEGs are suggested to include not only well-known OS in PD research candidates, but also novel candidates.

Moreover, PD-linked genes with limited evidentiary publications may harbor false positives (genes that are flagged as disease-associated in a database containing evidence with inappropriate evidence). For example, we identified superoxide dismutase 3 (*SOD3*) as a false-positive result. *SOD3* was linked to PD through the Open Targets Platform with a single literature annotation, which on closer inspection turns out that the contents in the literature claims the opposite that no significant SNP-PD risk association are found for *SOD*3^59^. This example demonstrates that our pipeline has the potential to reveal false positives entries in the database.

### Limitations

Finally, two limitations were identified in the analysis pipeline. First, several pipeline steps require time-consuming manual efforts. The curation of RNA-seq data is required for gene expression analysis in the first step. Following the refinement of the unexploited candidate genes, PMC was manually searched to assess the status of each gene. These manual steps render the methodology unsuitable for the comprehensively identification of numerous unexploited genes. Second, the referenced gene-disease databases are not static but will evolve with database research progress. We selected five databases to maximize the disease gene coverage presently. However, novel databases are likely to emerge and be integrated over time. Therefore, appropriate database selection based on contemporary availability is necessary.

## METHODS

### Curation of public data for OS

To collect OS RNA-sequencing data, relevant datasets were manually curated from the GEO^35^ repository based on five criteria: 1) total RNA or polyA-enriched sequencing, 2) samples under conditions related to the definition of OS, 3) samples under conditions related to increased ROS levels, 4) availability of paired normal-state samples as a control, and 5) cell cultures with brain relevance (neurons, astrocytes, and progenitors). This resulted in 122 matched OS-normal sample pairs for analysis as a result of curating started from August 2023 to October 31, 2023. Comprehensive details of the public datasets used in this study are listed in Supplementary Table 1^34^.

### Curation of public data for PD

To compile DEGs in PD, we conducted a manual survey searching the PubMed. Meta-analyses of three published studies were found and we extracted all reported DEGs. Additionally, genes queried for PD in the TWAS-Atlas database^39^ were incorporated into the PD-DEG list for downstream analysis. This curation was conducted from August 2023 to October 31, 2023. Comprehensive details regarding all the public datasets used in this study are listed in Table 2.

### OS meta-analysis

For RNA-seq data retrieval, processing, and quantification, we used Ikra^60^, an automated pipeline program for RNA-seq data of *Homo sapiens* and *Mus musculus*. The following pipeline comprised fasterq-dump (version.3.0.1)^61^, trim-galore (version.0.6.7)^62^, and salmon (version.1.4.0)^63^ processes, with reference transcript sets in GENCODE Release 44 (GRCh38.p14). The transcript IDs were converted into the gene IDs using tximport^64^ [Supplementary Data 2]^34^. To retrieve DEGs across 122 paired RNA-seq, we devised an oxidative stress-normal state score (ON-score) based on these datasets. Initially, the ON-ratio was calculated for each gene, representing the expression ratio between OS and normal states across all sample pairs (Equation 1). Subsequently, genes were then categorized as upregulated, downregulated, or unchanged based on the ON-ratio exceeding a ±1.5-fold threshold. Furthermore, the ON-score for each gene was calculated using Equation 2, which involved subtracting the number of downregulated samples from the number of upregulated samples. This scoring methodology was detailed extensively in a previous study^15^. The ON score measured how many of the 122 pairs of samples dysregulate the expression of each gene.

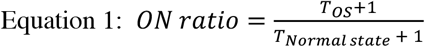

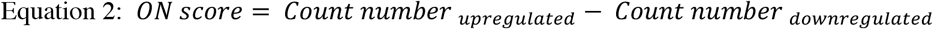

### Classifying genes by the availability of associations with PD

To classify genes based on prior evidence linking them to a disease, we originally developed the tool called gene-disease-linker^40^ [Figure 4]. This tool uses five databases (accessed on February 8, 2024) – Open Targets Platform^3^, RNAdisease^12^, miRTex^11^, DisGeNET^10^, and PubChem^13^ - to collect gene-disease relationships. By inputting a gene list into gene-disease-linker, it outputs whether there is a gene-disease association for each gene in the list regarding a specific disease. In cases a gene exhibiting an association with the disease, it also concurrently outputs the supporting literature with PubMed ID. In contract, no gene-disease association is indicated if there is no supporting literature annotated to a gene. In this study, gene-disease-linker was used to classify genes as either PD-linked (existing literature linking the gene to PD) or PD-unlinked (no evidence found) (executed on February 8, 2024). The source codes and usage of gene-disease-linker are available in the GitHub repository.

**Figure 4.**
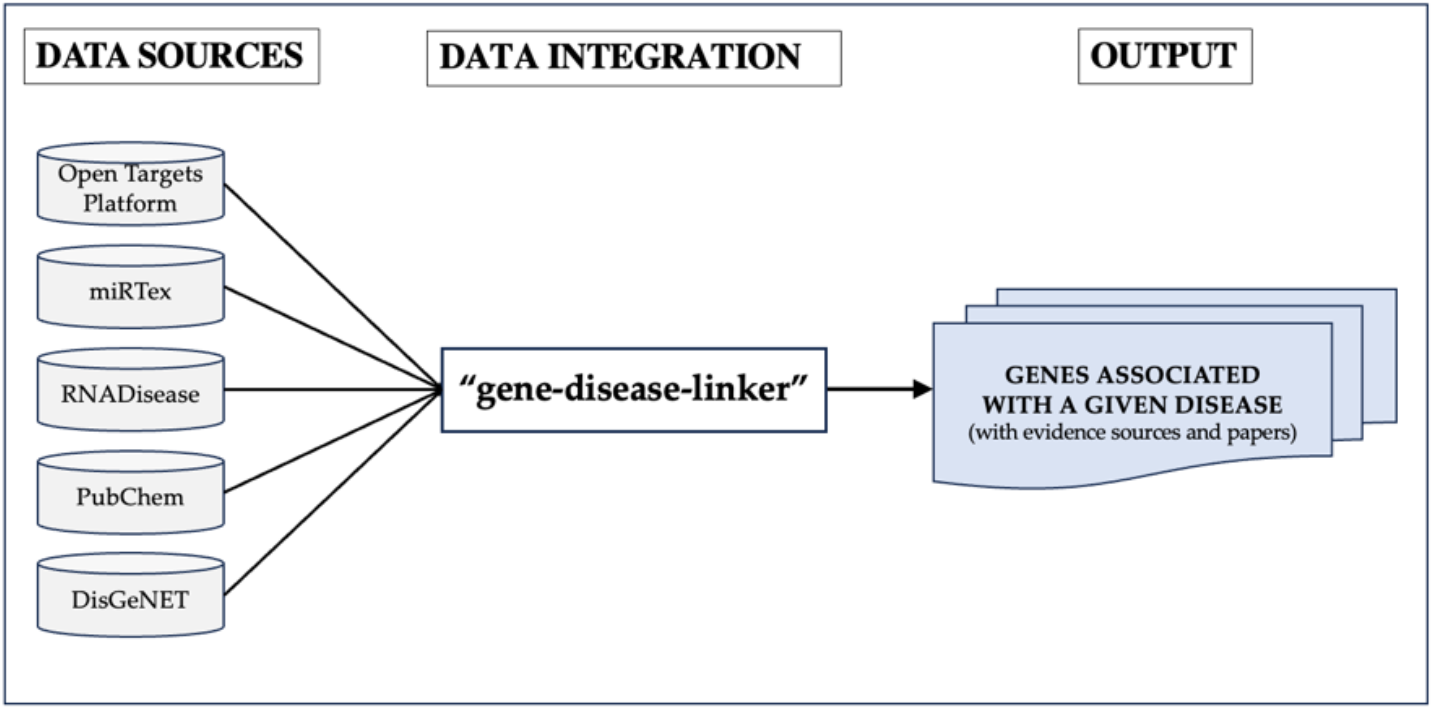
Overview of gene-disease-linker collecting information about gene-disease associations based on the relevant five databases.

### Criteria to judge PD unexploited genes

We searched gene names in PubMed Central, a full-text literature database with the following query: “GENE_NAME[All Fields] AND Parkinson[All Fields]”. Within the retrieved articles, we analyzed the surrounding textual context of gene mentions to identify descriptions indicating or suggesting molecular functional relationships with PD. Only genes with mechanistic evidence or relationships reported in the literature were judged unexploited. Genes only listed among the DEGs without any statements related to functional implications were excluded from unexploited.

### Other analysis

GSEA was performed using the web-based tool Metascape^65^. Shared genes among the various gene sets were visualized using a publicly available web-based Venn diagram generator^66^.

## DATA AVAILABILITY

The datasets curated, generated, and analyzed during this study are available in the figshare repository^34^.

## CODE AVAILABILITY

The underlying code for the current study is available at gene-disease-linker^40^.

### ACKNOWLEDGEMENTS

This research was supported by the Center of Innovation for Bio-Digital Transformation (BioDX), the open innovation platform for industry-academia co-creation (COI-NEXT), the Japan Science and Technology Agency (JST, COI-NEXT, JPMJPF2010), and the ROIS-DS-JOINT (007RP2023).

This work was also supported by the JST, which established university fellowships for the creation of science and technology innovation (Grant Number JPMJFS2129). Computations were performed on the computers at Hiroshima University Genome Editing Innovation Center. We also would like to thank all laboratory members at Hiroshima University and the Database Center of Life Science (DBCLS) for their valuable comments.

## AUTHORS INFORMATION

Authors and affiliations

Program of Biomedical Science, Graduate School of Integrated Sciences for Life, Hiroshima University, 3-10-23 Kagamiyama, Higashi-Hiroshima, Hiroshima 739-0046, Japan;

Takayuki Suzuki, Hidemasa Bono

Genome Editing Innovation Center, Hiroshima University, 3-10-23 Kagamiyama, Higashi-Hiroshima, Hiroshima 739-0046, Japan;

Hidemasa Bono

Database Center for Life Science (DBCLS), Joint Support-Center for Data Science Research, Research Organization of Information and Systems (ROIS), 178-4-4 Wakashiba, Kashiwa, Chiba 277-0871, Japan;

### Contributions

T.S. was responsible for data curation, software development, pipeline analysis, draft of the original manuscript. T.S and H. B were responsible for the study design, conceptualization, methodology manuscript review and editing. H.B. was responsible for the project administration, funding acquisition. All authors read and approved the final manuscript.

### Corresponding authors

Correspondence to Hidemasa Bono

## ETHICS DECLARATIONS

Competing interests

The authors declare no competing interests.

## Notes

https://doi.org/10.6084/m9.figshare.c.7114075.v1

### Competing Interest Statement

The authors have declared no competing interest.

## References

1. Rehm, H. L. et al. ClinGen — The Clinical Genome Resource. New England Journal of Medicine 372, 2235–2242 (2015).

2. Sollis, E. et al. The NHGRI-EBI GWAS Catalog: knowledgebase and deposition resource. Nucleic Acids Research 51, D977–D985 (2023).

3. Ochoa, D. et al. The next-generation Open Targets Platform: reimagined, redesigned, rebuilt. Nucleic Acids Research 51, D1353–D1359 (2023).

4. Tian, R. et al. Genome-wide CRISPRi/a screens in human neurons link lysosomal failure to ferroptosis. Nat Neurosci 24, 1020–1034 (2021).

5. Ghoussaini, M. et al. Open Targets Genetics: systematic identification of trait-associated genes using large-scale genetics and functional genomics. Nucleic Acids Research 49, D1311–D1320 (2021).

6. Dhindsa, R. S. et al. Rare variant associations with plasma protein levels in the UK Biobank. Nature 622, 339–347 (2023).

7. Karczewski, K. J. et al. Systematic single-variant and gene-based association testing of thousands of phenotypes in 394,841 UK Biobank exomes. Cell Genomics 2, 100168 (2022).

8. The Europe PMC Consortium. Europe PMC: a full-text literature database for the life sciences and platform for innovation. Nucleic Acids Research 43, D1042–D1048 (2015).

9. Kafkas, Ş., Dunham, I. & McEntyre, J. Literature evidence in open targets - a target validation platform. Journal of Biomedical Semantics 8, 20 (2017).

10. Piñero, J. et al. DisGeNET: a comprehensive platform integrating information on human disease-associated genes and variants. Nucleic Acids Research 45, D833–D839 (2017).

11. Li, G. et al. miRTex: A Text Mining System for miRNA-Gene Relation Extraction. PLOS Computational Biology 11, e1004391 (2015).

12. Chen, J. et al. RNADisease v4.0: an updated resource of RNA-associated diseases, providing RNA-disease analysis, enrichment and prediction. Nucleic Acids Research 51, D1397–D1404 (2023).

13. Li, Q., Kim, S., Zaslavsky, L., Cheng, T. & Yu, B. Resource Description Framework (RDF) Modeling of Named Entity Co-occurrences Derived from Biomedical Literature in the PubChemRDF.

14. Esaki, T. & Ikeda, K. Difficulties and prospects of data curation for ADME in silico modeling. CBIJ 23, 1–6 (2023).

15. Suzuki, T., Ono, Y. & Bono, H. Comparison of Oxidative and Hypoxic Stress Responsive Genes from Meta-Analysis of Public Transcriptomes. Biomedicines 9, 1830 (2021).

16. Bono, H. Meta-Analysis of Oxidative Transcriptomes in Insects. Antioxidants 10, 345 (2021).

17. Dorsey, E. R. et al. Global, regional, and national burden of Parkinson’s disease, 1990–2016: a systematic analysis for the Global Burden of Disease Study 2016. The Lancet Neurology 17, 939–953 (2018).

18. Davie, C. A. A review of Parkinson’s disease. British Medical Bulletin 86, 109–127 (2008).

19. Obeso, J. A. et al. Missing pieces in the Parkinson’s disease puzzle. Nat Med 16, 653–661 (2010).

20. Wiecki, T. V. & Frank, M. J. Chapter 14 - Neurocomputational models of motor and cognitive deficits in Parkinson’s disease. in Progress in Brain Research (eds. Björklund, A. & Cenci, M. A.) vol. 183 275–297 (Elsevier, 2010).

21. Sahoo, S., Padhy, A. A., Kumari, V. & Mishra, P. Role of Ubiquitin–Proteasome and Autophagy-Lysosome Pathways in α-Synuclein Aggregate Clearance. Mol Neurobiol 59, 5379–5407 (2022).

22. Zhou, Z. D., Yi, L. X., Wang, D. Q., Lim, T. M. & Tan, E. K. Role of dopamine in the pathophysiology of Parkinson’s disease. Translational Neurodegeneration 12, 44 (2023).

23. Ramesh, S., Arachchige, A. S. P. M., Ramesh, S. & Arachchige, A. S. P. M. Depletion of dopamine in Parkinson’s disease and relevant therapeutic options: A review of the literature. AIMSN 10, 200–231 (2023).

24. Dias, V., Junn, E. & Mouradian, M. M. The Role of Oxidative Stress in Parkinson’s Disease. Journal of Parkinson’s Disease 3, 461–491 (2013).

25. Klinkovskij, A., Shepelev, M., Isaakyan, Y., Aniskin, D. & Ulasov, I. Advances of Genome Editing with CRISPR/Cas9 in Neurodegeneration: The Right Path towards Therapy. Biomedicines 11, 3333 (2023).

26. Simmnacher, K. et al. Unique signatures of stress-induced senescent human astrocytes. Experimental Neurology 334, 113466 (2020).

27. Krauskopf, J. et al. Transcriptomics analysis of human iPSC-derived dopaminergic neurons reveals a novel model for sporadic Parkinson’s disease. Mol Psychiatry 27, 4355–4367 (2022).

28. Tong, Z.-B., Braisted, J.Chu, P.-H. & Gerhold, D. The MT1G Gene in LUHMES Neurons Is a Sensitive Biomarker of Neurotoxicity. Neurotox Res 38, 967–978 (2020).

29. The Irradiated Brain Microenvironment Supports Glioma Stemness and Survival via Astrocyte-Derived Transglutaminase 2 | Cancer Research | American Association for Cancer Research. https://aacrjournals.org/cancerres/article/81/8/2101/670586/The-Irradiated-Brain-Microenvironment-Supports.

30. Shimada, M., Tsukada, K., Kagawa, N. & Matsumoto, Y. Reprogramming and differentiation-dependent transcriptional alteration of DNA damage response and apoptosis genes in human induced pluripotent stem cells. Journal of Radiation Research 60, 719–728 (2019).

31. Loeliger, B. W. et al. Effect of Ionizing Radiation on Transcriptome during Neural Differentiation of Human Embryonic Stem Cells. rare 193, 460–470 (2020).

32. Murotomi, K. et al. Cyclo-glycylproline attenuates hydrogen peroxide-induced cellular damage mediated by the MDM2-p53 pathway in human neural stem cells. Journal of Cellular Physiology 238, 434–446 (2023).

33. Crowe, E. P. et al. Changes in the Transcriptome of Human Astrocytes Accompanying Oxidative Stress-Induced Senescence. Frontiers in Aging Neuroscience 8, (2016).

34. Suzuki, T. A systematic exploration of unexploited disease-related genes. (2024) doi:10.6084/m9.figshare.c.7114075.v1.

35. Barrett, T. et al. NCBI GEO: archive for functional genomics data sets—update. Nucleic Acids Research 41, D991–D995 (2013).

36. Mariani, E. et al. Meta-Analysis of Parkinson’s Disease Transcriptome Data Using TRAM Software: Whole Substantia Nigra Tissue and Single Dopamine Neuron Differential Gene Expression. PLOS ONE 11, e0161567 (2016).

37. Phung, D. M. et al. Meta-Analysis of Differentially Expressed Genes in the Substantia Nigra in Parkinson’s Disease Supports Phenotype-Specific Transcriptome Changes. Frontiers in Neuroscience 14, (2020).

38. Cappelletti, C. et al. Transcriptomic profiling of Parkinson’s disease brains reveals disease stage specific gene expression changes. Acta Neuropathol 146, 227–244 (2023).

39. Lu, M. et al. TWAS Atlas: a curated knowledgebase of transcriptome-wide association studies. Nucleic Acids Research 51, D1179–D1187 (2023).

40. szktkyk. szktkyk/gene-disease-linker. (2024).

41. Index of /gene/DATA. https://ftp.ncbi.nlm.nih.gov/gene/DATA/.

42. Kia, D. A. et al. Identification of Candidate Parkinson Disease Genes by Integrating Genome-Wide Association Study, Expression, and Epigenetic Data Sets. JAMA Neurology 78, 464–472 (2021).

43. Li, Y. I., Wong, G., Humphrey, J. & Raj, T. Prioritizing Parkinson’s disease genes using population-scale transcriptomic data. Nat Commun 10, 994 (2019).

44. Chen, L. et al. Study of molecular patterns associated with ferroptosis in Parkinson’s disease and its immune signature. PLOS ONE 18, e0295699 (2023).

45. Chung, S.-K. & Lee, S.-Y. Advances in Gene Therapy Techniques to Treat LRRK2 Gene Mutation. Biomolecules 12, 1814 (2022).

46. Liu, J. et al. NUPR1 is a critical repressor of ferroptosis. Nat Commun 12, 647 (2021).

47. Wang, X. et al. UHRF2 regulates cell cycle, epigenetics and gene expression to control the timing of retinal progenitor and ganglion cell differentiation. Development 149, dev195644 (2022).

48. Bucher, M. L. et al. Acquired dysregulation of dopamine homeostasis reproduces features of Parkinson’s disease. npj Parkinsons Dis. 6, 1–13 (2020).

49. Choi, W.-S.Kim, H.-W. & Xia, Z. JNK inhibition of VMAT2 contributes to rotenone-induced oxidative stress and dopamine neuron death. Toxicology 328, 75–81 (2015).

50. Su, C.-J. et al. Thioredoxin-interacting protein induced α-synuclein accumulation via inhibition of autophagic flux: Implications for Parkinson’s disease. CNS Neuroscience & Therapeutics 23, 717–723 (2017).

51. Yang, D. et al. Neurofilament light chain as a mediator between LRRK2 mutation and dementia in Parkinson’s disease. npj Parkinsons Dis. 9, 1–6 (2023).

52. Gong, L. et al. Neurofilament Light Chain (NF-L) Stimulates Lipid Peroxidation to Neuronal Membrane through Microglia-Derived Ferritin Heavy Chain (FTH) Secretion. Oxidative Medicine and Cellular Longevity 2022, e3938940 (2022).

53. Gellhaar, S., Sunnemark, D., Eriksson, H., Olson, L. & Galter, D. Myeloperoxidase-immunoreactive cells are significantly increased in brain areas affected by neurodegeneration in Parkinson’s and Alzheimer’s disease. Cell Tissue Res 369, 445–454 (2017).

54. Maki, R. A. et al. Human myeloperoxidase (hMPO) is expressed in neurons in the substantia nigra in Parkinson’s disease and in the hMPO-α-synuclein-A53T mouse model, correlating with increased nitration and aggregation of α-synuclein and exacerbation of motor impairment. Free Radical Biology and Medicine 141, 115–140 (2019).

55. Verdiperstat | ALZFORUM. https://www.alzforum.org/therapeutics/verdiperstat.

56. Chang, C. Y., Choi, D.-K., Lee, D. K., Hong, Y. J. & Park, E. J. Resveratrol Confers Protection against Rotenone-Induced Neurotoxicity by Modulating Myeloperoxidase Levels in Glial Cells. PLOS ONE 8, e60654 (2013).

57. Zhao, J. et al. LINC00938 alleviates hypoxia ischemia encephalopathy induced neonatal brain injury by regulating oxidative stress and inhibiting JNK/p38 MAPK signaling pathway. Experimental Neurology 367, 114449 (2023).

58. Yousefi, M., Peymani, M., Ghaedi, K., Irani, S. & Etemadifar, M. Significant modulations of linc001128 and linc0938 with miR-24-3p and miR-30c-5p in Parkinson disease. Sci Rep 12, 2569 (2022).

59. Liu, C., Fang, J. & Liu, W. Superoxide dismutase coding of gene polymorphisms associated with susceptibility to Parkinson’s disease. JIN 18, 299–303 (2019).

60. Yu, H. et al. yyoshiaki/ikra: ikra v2.0.1. Zenodo https://doi.org/10.5281/zenodo.5541399 (2021).

61. The NCBI SRA (Sequence Read Archive); NCBI—National Center for Biotechnology Information/NLM/NIH: Bethesda, MD, USA, 2021.

62. Babraham Bioinformatics - Trim Galore! https://www.bioinformatics.babraham.ac.uk/projects/trim_galore/.

63. Patro, R., Duggal, G., Love, M. I., Irizarry, R. A. & Kingsford, C. Salmon provides fast and bias-aware quantification of transcript expression. Nat Methods 14, 417–419 (2017).

64. tximport. Bioconductor http://bioconductor.org/packages/tximport/.

65. Zhou, Y. et al. Metascape provides a biologist-oriented resource for the analysis of systems-level datasets. Nat Commun 10, 1523 (2019).

66. Draw Venn Diagram. https://bioinformatics.psb.ugent.be/webtools/Venn/.

